# Classification of mouse ultrasonic vocalizations using deep learning

**DOI:** 10.1101/358143

**Authors:** A. Ivanenko, P. Watkins, M. A. J. van Gerven, K. Hammerschmidt, B. Englitz

## Abstract

Vocalizations are a widespread means of communication in the animal kingdom. Mice use a large repertoire of ultrasonic vocalizations (USVs) in different social contexts, for instance courtship, territorial dispute, dominance and mother-pup interaction. Previous studies have pointed to differences in the USVs in different context, sexes, strains and individuals, however, in many cases the outcomes of the analyses remained inconclusive.

We here provide a more general approach to automatically classify USVs using deep neural networks (DNN). We classified the sex of the emitting mouse (C57Bl/6) based on the vocalization’s spectrogram, reaching unprecedented performance (~84% correct) in comparison with other techniques (Support Vector Machines: 64%, Ridge regression: 52%). Vocalization characteristics of individual mice only contribute mildly, and sex-only classification reaches ~78%. The performance can only partially be explained by a set of classical shape features, with duration, volume and bandwidth being the most useful predictors. Splitting estimation into two DNNs, from spectrograms to features (57-82%) and features to sex (67%) does not reach the single-step performance.

In summary, the emitter’s sex can be successfully predicted from their spectrograms using DNNs, excelling over other classification techniques. In contrast to previous research, this suggests that male and female vocalizations differ in their spectrotemporal structure, recognizable even in single vocalizations.

## Introduction

Identification of sex on the basis of sensory cues provides important information for successful reproduction. When listening to a conversation, humans can typically make an educated guess about the sexes of the participants. Limited research on this topic has identified multiple acoustic predictors, ranging from the fundamental frequency to formant measures (Pisanski et al. 2016).

Similar to humans, mice vocalize in particular during social interactions (Chabout et al. 2015; Heckman et al. 2016; Heckman et al. 2017; Neunuebel et al. 2015; Portfors and Perkel 2014). The complexity of the vocalizations produced during social interactions can be substantial (Holy and Guo 2005). While in humans and other species sex-specific differences in body dimensions (vocal tract length, vocal fold characteristics) lead to predictable differences in vocalization (Markova et al. 2016; Pfefferle and Fischer 2006), the vocal tract properties of male and female have not been shown to differ significantly (Mahrt et al. 2016; Roberts 1975). Hence, for mice the expected differences in male/female USVs are less predictable from physiological characteristics.

Previous research on the properties of male and female ultrasonic vocalizations (USVs) (Hammerschmidt et al. 2012) found differences in usage of vocal types, but could not identify reliable predictors (of the emitter’s sex) on the basis of single vocalizations. Yet, experiments playing male mouse courtship songs to adult females suggest that at least females are able to guess the sex of other mice based on single vocalizations (Hammerschmidt et al. 2009; Shepard and Liu 2011)

In the present study, we revisit this question by making use of advanced machine learning techniques, in particular deep learning (LeCun et al. 2015). We estimate classifiers for sex of the emitter on the basis of the vocalizations spectrogram or an intermediate level of acoustic properties using a range of different approaches. Vocalizations from one of the most common laboratory strains, C57Bl/6 were used, in which one of the mice was anesthetized, thus uniquely associating a sex with each vocalization (Hammerschmidt et al. 2012)

We find that spectrogram-based classification using a deep neural network reaches an average accuracy of 84%, which is substantially greater than previous approaches as well as presently tested linear (ridge regression) or nonlinear (support vector machines, SVM, (Cortes and Vapnik 1995)(classifiers. This performance can partially be explained by statistical differences in particular features between male and female vocalizations. While individually, these differences are insufficient for classification due to a high degree of variability, they are nonlinearly combined by the DNN to achieve this level of classification. We demonstrate that this combination has to already occur on the level of the spectrogram, since a classification on the basis of a conventional feature set does not achieve similar levels of performance (67%). We further provide a network that can automatically classify the properties, and thus automatizes an otherwise a time-consuming, manual task.

In summary, the ability to classify the sex based on vocalizations alone provides an important building-block for analyzing vocalizations during social interaction. The present results indicate that mouse vocalizations can typically be attributed to male or female mice, however, the relation to the sex-specific spectrotemporal properties of the USVs remains to be fully addressed.

## Results

We reanalyzed recordings of ultrasonic vocalizations (USVs) from single mice during a social interaction with an anesthetized mouse (N=17, in 9 female awake, in 8 male awake, Fig. 1A, (Hammerschmidt et al. 2012)). The awake mouse vocalized actively in each recording (Male: 181±32 min^−1^, Female: 212±14 min^−1^, Fig. 1B) giving a total of 10055 (female: 5723, male: 4332) automatically extracted USVs of varying spectrotemporal structure and uniquely known sex of the emitter (Fig. 1C). Previous approaches for assessing the gender using basic analysis have not lead to single-vocalization level predictability (Hammerschmidt et al. 2012). Here, we utilized the powerful and general framework of deep neural networks (DNNs) to predict the emitter’s sex for single vocalizations (Fig. 1D). Obtaining best-in-class performance on this challenge we further investigate the basis for this performance. For this purpose, separate DNNs were trained for predicting features from spectrograms as well as sex from features, on the basis of human classified features.

**Figure 1:**
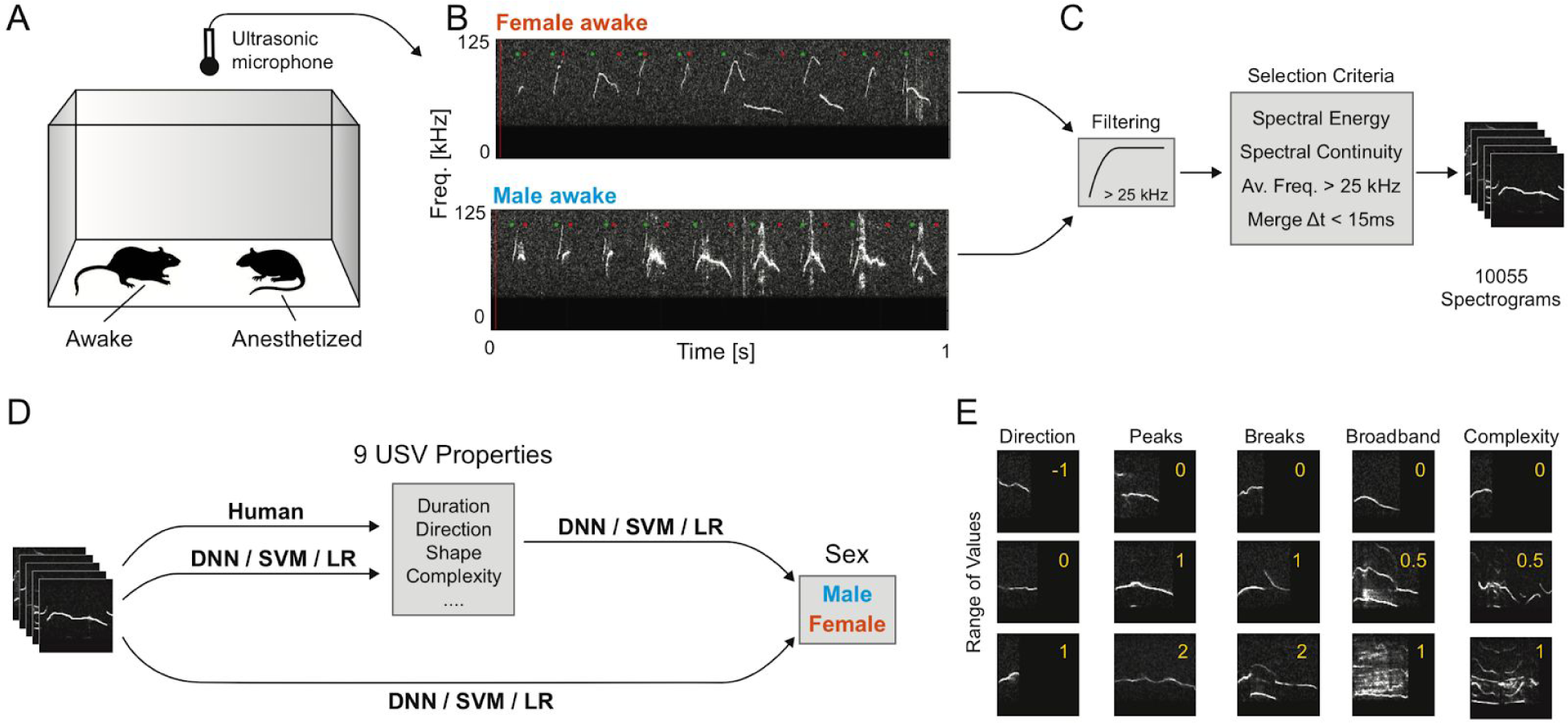
Recording and Classifying Mouse Vocalizations. **A** Mouse vocalization were recorded from a pair of mice, in which one was awake, while the other was anesthetized, allowing an unambiguous attribution of the recorded vocalizations. **B** Vocalization from male and female mice (recorded in separate sessions) share a lot of properties, while differing in others. The present samples were picked at random and indicate that differences exist, while other samples would look more similar. **C** Vocalizations were automatically segmented using a set of filtering and selection criteria (see Methods for details), leading to a total set of 10055 vocalizations. **D** We aimed to estimate the properties and the sex of its emitter for individual vocalizations. First, the ground truth for the properties were established by a human classifier. We next estimated 3 relations, Spectrogram-Properties, Properties-Sex and Spectrogram-Sex directly, using both a Deep Neural Network (DNN), support vector machines (SVM) and regularized linear regression (LR). **E** The properties attributed manually to individual vocalizations could take different values (rows, red number in each subpanel), illustrated here for a subset of the properties (columns). See Methods for a detailed list and description of the properties.

*Basic USV features differ between the sexes, but are insufficient to distinguish single USVs* While previous approaches have indicated few differences between male and female vocalizations ((Hammerschmidt et al. 2012), but see (Heckman et al. 2017) for CBA mice), we reassess this question first using a set of 9 hand-picked features, quantifying spectrotemporal properties of the USVs (see Fig. 1E and Method for details). The first three were quantified automatically, while the latter 6 were scored by human classifiers.

We find significant differences in multiple features (7/9, see Fig. 2A-I for details, based on Wilcoxon Signed Ranks test) between the sexes (in all figures: red = female, blue = male). This suggests that there is exploitable information about the emitter’s sex. However, the substantial variability across USVs renders each feature in isolation insufficient for classifying single USVs (see percentiles in Fig. 2A-I). Next, we investigated whether the joint set of features or the raw spectrograms have some sex-specific properties that can be exploited for classification.

**Figure 2:**
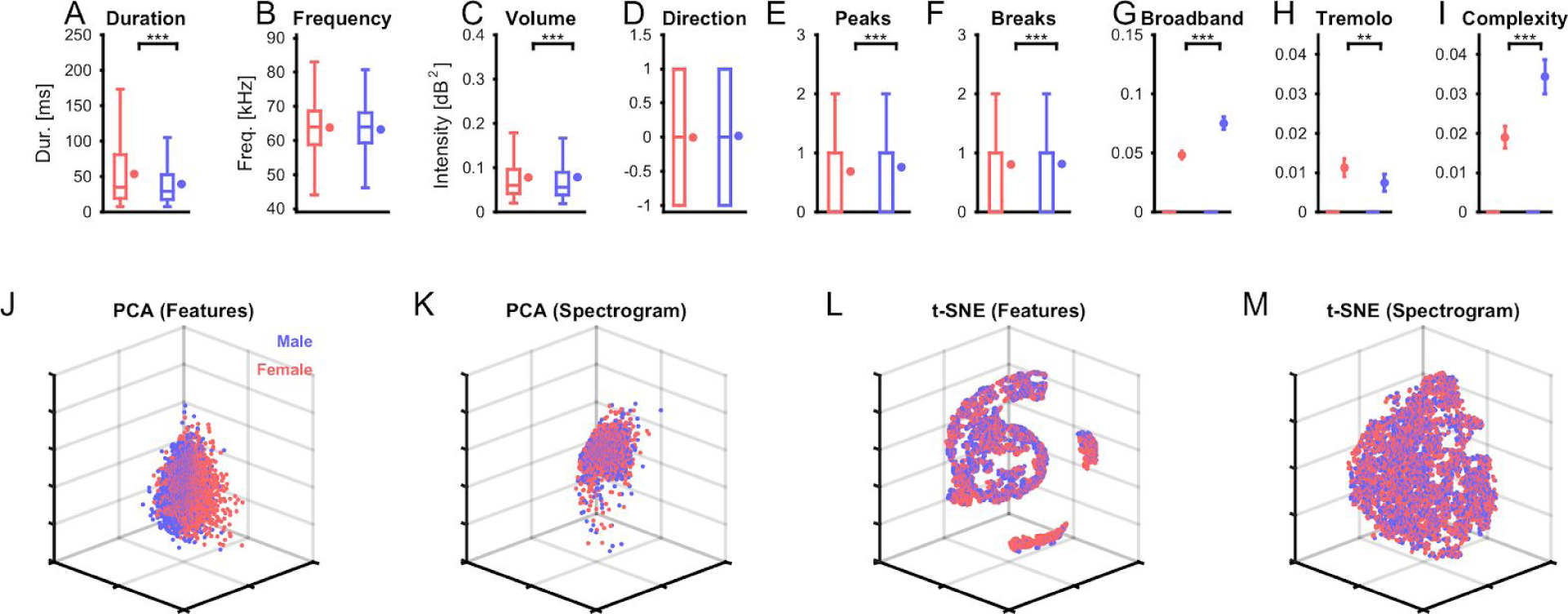
Basic Sex-Dependent Differences between Vocalizations. (**A-I**) We quantified a range of properties for single vocalizations (see Methods for details) and compared them across the sexes (blue: male, red: female). Most properties exhibited significant differences in median between the sexes (Wilcoxon rank sum test), except for the average frequency (**B**) and directionality (**D**). However, given the distribution of the data (box-plots, left in each panel), the variability across each property nonetheless makes it hard to use individual properties of determining the sex of the emitter. The graphs on the right for each color in each panel, show the mean and SEM. In **G-I**, only few USVs has values different than 0, hence the box-plots are sitting at 0. Outliers are not shown to avoid crowding the plots. (**J-M**) Dimensionality reduction can reveal more complex relationships between multiple properties. We computed principal components (PCA) and t-statistic stochastic neighborhood embedding (t-SNE) for both the features (**J/L**) and the spectrograms (**K/M**). In particular, feature-based t-SNE (**L**) obtained some interesting groupings, which did, however, not separate between the sexes (red vs. blue). Each dot represents a single vocalization, after dimensionality reduction. Axes are left unlabelled, since they represent a combination of properties.

Applying principal component analysis (PCA) to the 9-dimensional set of features and retaining three dimensions, a largely intermingled representation is obtained (Fig. 2J). PCA of the spectrograms provides a slightly more structured spatial layout in the most variable three dimensions (Fig. 2K). However, as for the features, the sexes overlap in space with no apparent separation. The basic dimensionality reduction performed by PCA, hence, reflects the co-distribution of properties and spectrograms between the sexes. At the same time it fails to even pick-up the basic properties along which the sexes differ (see above).

Using instead t-Distributed Stochastic Neighborhood Embedding (tSNE, (Maaten and Hinton 2008)(for dimensionality reduction, reveals a clustered structure for both features (Fig. 2L) and spectrograms (Fig. 2M). For the features the clustering is very clean, and reflects the different values of the features (e.g. different number of breaks define different clusters, or gradients within clusters (Duration/Frequency), not shown here, since the focus is on sex classification). However, USVs from different sexes are quite close in space and co-occur in the all clusters, although density-differences (of male or female USVs) exist in individual clusters. For the spectrograms the representation is less clear, but still shows much clearer clustering than after PCA. Mapping feature properties to the local clusters did not lead to an intelligible separation (data not shown).

In summary, spectrotemporal features in isolation or in basic conjunction appear insufficient to reliably classify single vocalizations by their emitter’s sex. However, the prior analyses do not attempt to directly learn sex-specific spectrotemporal structure from the given data-set, but instead assume a restricted set of features. Next, we used data-driven classification algorithms to directly learn differences between male and female vocalizations.

### Convolutional deep neural network identifies the emitter’s sex from single USVs

Akin to the recent successes of convolutional deep neural networks (cDNNs) in other fields, e.g. machine vision (LeCun et al. 2015), we focus here on a direct classification of the emitter’s sex based on the spectrograms of single USVs. The architecture chosen is a - by now - classical network structure of convolutional layers, followed by fully connected layers, and a single output representing the probability of a male or female source (see Methods for details and Fig. 3A). The cDNN performs surprisingly well, in comparison to other classification techniques (see below).

**Figure 3:**
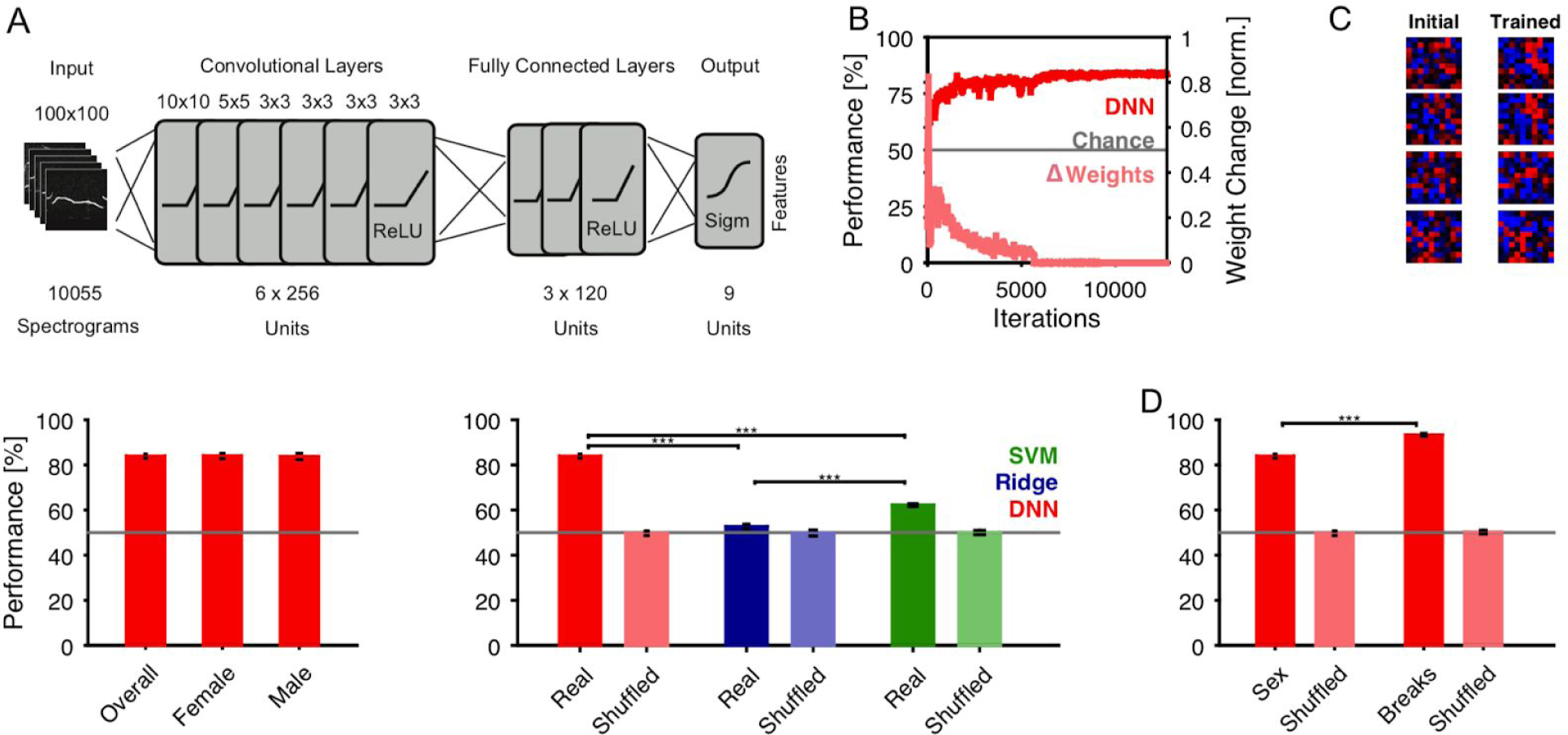
Deep neural network reliably determines the emitter’s sex from individual vocalizations. **A** We trained a deep neural network (DNN) with 6 convolutional and 3 fully connected layers to classify the sex of emitter from the spectrogram of the vocalization. **B** The network’s performance on the test set rapidly improved to an asymptote of ~85% (dark red), clearly exceeding chance. Correspondingly the change in the network’s weights (light red) progressively decreased after stabilizing after ~6k iterations. Data shown for a representative training run. **C** The shape of the input fields in the first convolutional layer became more reminiscent of the tuning curves in auditory system (Ahrens et al. 2008; Englitz et al. 2010). Samples are representatively chosen among the entire set of 256 units in this layer. **D** The average performance of the DNN (10 runs of crossvalidation) was 83.9±0.9%, which did not differ significantly between male and female vocalizations (p>0.05, Wilcoxon rank sum test). **E** The DNN performance by far exceeded the performance of ridge regression (regularized linear regression, blue, 52.7±1.0%) and support vector machines (SVM, green, 62.2±0.6%). Bars in light colors show the corresponding estimation with randomized labels, which are all at chance level (gray line) **F** The performance by the DNN was not only limited by the properties of the spectrograms (e.g. background noise, sampling, etc.) since a DNN trained on the number of breaks (right bars) performed significantly better. This control shows that the identical set of stimuli can be better separated on a simpler (but also binary) task. Light bars again show performance on randomized labels.

During the training procedure the cDNN’s parameters are adapted to optimize performance on the test set, which was not used for training (see Methods for details on crossvalidation). Performance starts out near chance level (red, Fig. 3B), with the initial iterations leading to substantial changes in weights (light red). It takes ~6k iterations before the cDNN settles close to its final, asymptotic performance (here 85%, see below for average).

The learning process modifies the properties of the DNN throughout all layers. In the first layer, the neurons parameters can be visualized in the stimulus domain, i.e. similar to spatial receptive fields for real neurons. Initial random configurations adapt to become more classical local filters, e.g. shaped like local patches or lines (Fig. 3C). This behavior is well documented for visual classification tasks, however, it is important to verify it for the current, limited set of auditory stimuli.

The classification performance, averaged over 10 crossvalidation runs, was stable around 83.9±0.9% (mean and SEM across vocalizations, Fig. 3D). Female classification was slightly more accurate in comparison to male USVs (p<0.05, Wilcoxon signed ranks test).

In comparison with more classical classification techniques, such as regularized linear regression (Ridge, blue, Fig. 3E) and support vector machines (SVM, green), the DNN (red) performs significantly better (Ridge: 52.7±1.0%; SVM 62.2±0.6%; DNN: 83.9±0.9%; all comparisons: p<0.001). Shuffling the sexes of all vocalizations leads to chance performance, and thus indicates that the performance is not based on overlearning properties of a random set (Fig. 3E, light colors). Instead it has to rely on the characteristic property within one sex (see below for contributions by individual animals).

Lastly, to verify that the general properties of the spectrograms do not pose a sex-independent limit on classification performance, we evaluated the performance of the same network on another complex task: to determine whether a vocalization has a break or not (this is generalized in Fig. 5 below to multiple breaks). On the latter task the network can do significantly better than on classifying the emitter’s sex, reaching 93.3% (p<0.001, Wilcoxon signed ranks test, Fig. 3F).

Altogether, the estimated cDNN clearly outperforms classical competitors on the task and reaches a substantial performance, which could potentially be improved, e.g. by the addition of more data for refining the DNN’s parameters.

### Classification performance of sex generalizes across mice

DNNs are powerful classification devices with many thousands of parameters (LeCun et al. 1999) Despite techniques for generalizing their performance, they are in principle capable of learning to classify a large number of categories (Krizhevsky et al. 2017) In the present case, the performance of the above network could thus be based on the individual differences in vocalizations between mice, rather than common properties due to their sex. To exclude this possibility, we first verify that the performance is not just based on a few individuals (Fig. 4A), and secondly, prevent the network from learning individuals during the training phase (Fig. 4B). In the following section, we directly train on individuals.

**Figure 4:**
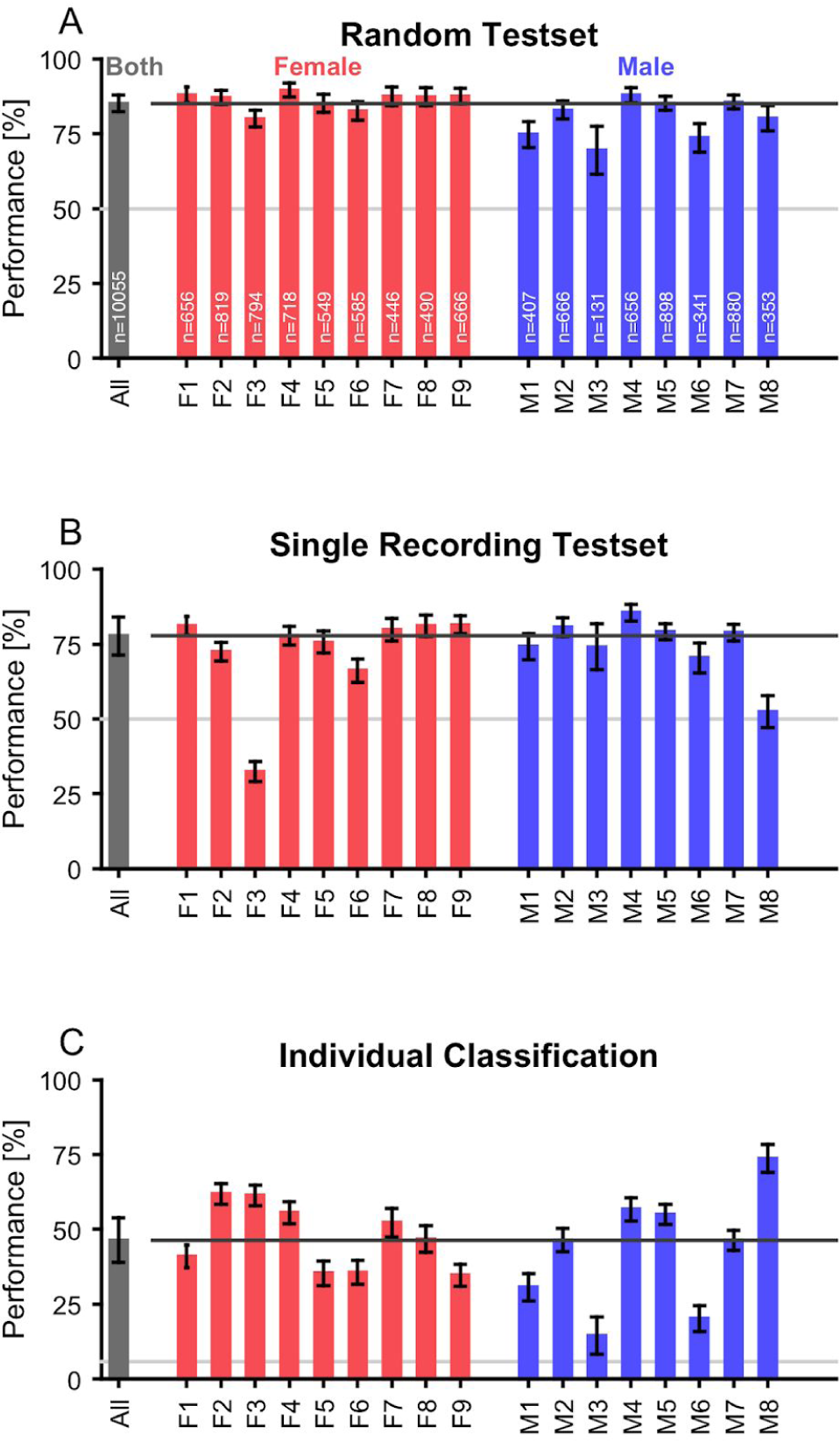
The sex classification is based on a mixture of individual and general properties. **A** Separating the sex classification results by individual mice demonstrates that the average performance is representative. The number of vocalizations is indicated in **A** for each individual. Coincidentally, the number of vocalizations emitted by 2 males and females are matched (verified to be correct). The light gray line indicates chance, and the dark gray line average performance. **B** We verified that the network does not just learn individual mice, by training on N-1 recordings, and using the remaining recording as the test set. Prediction performance remained comparatively high for individual mice, even increased for most males, with two exceptions. In this test configuration, the network cannot use properties of individual mice for prediction, but has to rely exclusively on properties general to a given sex. **C** We trained another DNN to classify the mouse identity, achieving above chance performance for almost all mice, indicating that differences between individual mice can also contribute to classification of sex. For classification of individuals, chance level is at 100/N_mice_ %.

For verifying that the performance is uniform across individuals, we perform a post-hoc separation of the network performance across individual mice (Fig. 4A, 85.1±2.9% median across mice). In particular for the female animals the performance is extremely homogeneous (red), while it shows some more variability for the male animals (blue), however, for all animals above chance.

To prevent the network from using individual animals, we reran the training, however, also using a full recording (corresponding to a single mouse) as the test set. Hence, during training the cDNN has no information about the mouse that it eventually predicts, beyond regularities based on the sex. The performance is reduced (77.7±6.3%, Fig. 4B), however, this is mainly driven by two animals (F3, M8). Hence, classification of sex is truly captured by the DNN. The discrepancy in performance to the random test may be explained by individual differences between mice, which are unavailable for classification if a mouse is excluded entirely from the training set.

### Classification of individual mice is possible but not necessary for sex classification

While the above indicates that individual features are not necessary for predicting the emitter’s sex, they may still contribute. This was addressed directly, by training another cDNN on the spectrograms, but predicting mouse identity instead of sex (see Methods for details on the output). The average performance is 46.4±7.5% (median across animals, Fig. 4C), i.e. far above chance performance (5.9% if sex is unknown, or at most 12.5% if the sex is known), which is also reflected in all of the individual animals (M3 has the largest p-value at 0.0002, binomial test against chance level). Hence, while not fully identifiable, individual mice appear to shape their vocalizations in characteristic ways, which is likely used in parts by our main network for sex-classification (Fig. 3), explaining the difference in performance for animal F3 between random (Fig. 4A) and single recording (Fig. 4B) test sets.

### USV features are insufficient to explain cDNN performance on sex classification

Simple combinations of features were insufficient to identify the emitter’s sex from individual USVs, as demonstrated above (Fig. 2). However, it could be that more complex combinations of the same features would be sufficient to reach similar performance as for the spectrogram-based classification (Fig. 3). We address this question by predicting the sex from the features alone - as classified by a human - using another DNN (see Methods for details and below). Further, we check whether the human-classified features can be learned by a cDNN. Together, these two steps provide a stepwise classification of sex from spectrograms.

First, we estimated separate cDNNs for 4 of the 6 features, i.e. direction, number of breaks, number of peaks and spectral breadth of activation (‘broadband’). The remaining two features - tremolo and complexity - were omitted, since most vocalizations scored very low in these values, thus creating a very skewed training set of the networks. A near optimal - but uninteresting - learning outcome is then a flat, near zero classification. The network structure of these cDNNs was chosen identical to the sex-classification network, for simpler comparison (Fig. 5A).

**Figure 5:**
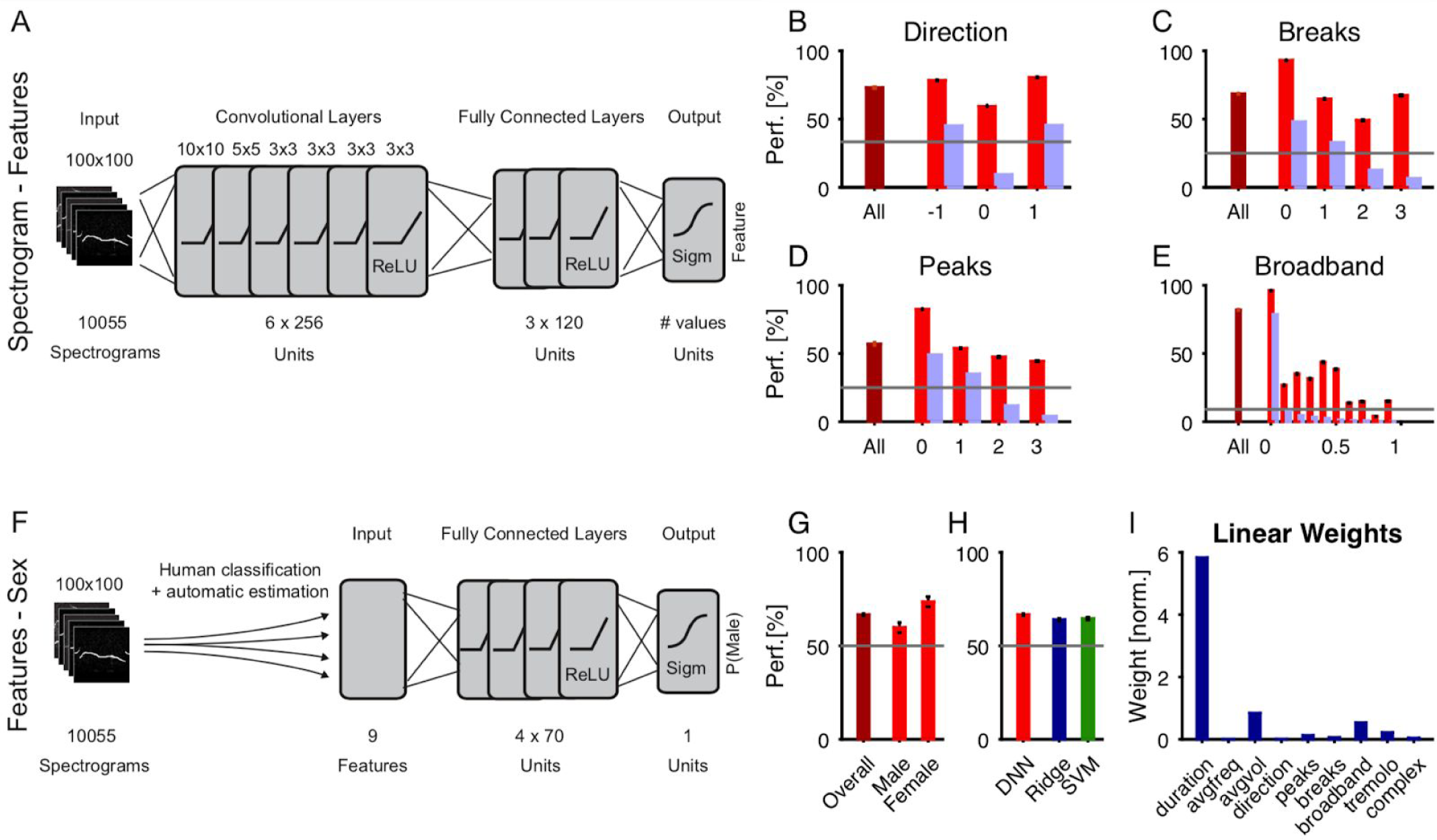
Features alone are insufficient to explain the DNN classification performance. **A** Features of individual vocalizations can also be measured using dedicated convolutional DNNs, one per feature, with identical architecture as for sex classification (see Fig. 3A). **B-E** Classification performance for different properties was robust, ranging between 57.0 and 82.0% on average (maroon) and depending on the individual value of each property (red). We trained networks for direction ({-1,0,1}, **B**), the number of breaks ({0-3}, **C**), the number of peaks ({0-3}, **D**) and the degree of broadband activation ([0,1], **E**). For the other 2 properties (complex and tremolo), most values were close to 0 and thus networks did not have sufficient training data for the other values. The light gray lines indicate chance performance, which depends on the number of choices for each property. The light blue bars indicate the distributions of values, also in %. **F** Using a non-convolutional DNN, we investigated how predictable features alone would be, i.e. without any information about the precise spectral structure of each vocalization. **G** Prediction performance was above chance (maroon, 66.7±0.5%) but far less than the prediction of sex on the basis of the raw spectrograms (see Fig. 3). Female USVs could be predicted more accurately than male USVs (see main text for statistics). Gray line indicates chance performance **H** Feature-based prediction of sex with DNNs was only slightly, but significantly better than using either ridge regression (linear, blue) or SVMs (red, see main text for statistics). **I** Duration, volume and the level of broadband activation were the most significant linear predictors for sex, when using ridge regression.

Average performance for all four features was far above chance (Fig. 5B-E, maroon, respective chance levels shown in gray), with some variability across the different values of each property (Fig. 5B-E, red). This variability is likely a consequence ofthe distribution of values in the overall set of USVs (Fig. 5B-E, light blue, scaled also in percentage) in combination with their inherent difficulty in prediction as well as higher variability in human assessment. Except for the broadband classification (82.0±0.4%), the classification performance for the features (Direction: 73.0±0.5%; Breaks: 68.6±0.4%; Peaks: 57.0±1.0%) stayed below the one for sex (83.9±0.9%).

Second, we estimated a DNN without convolutional layers for predicting the emitter’s sex from the chosen 9 features (see Methods for details, and Fig. 5F). The overall performance of the DNN was well above chance (66.7±0.5%, Fig. 5G, maroon), but remained even further below the raw spectrogram performance (83.9%) than the feature classification. Female vocalizations were recognized significantly more frequently (Fig. 5G, red). In comparison with ridge regression (64.0±0.7%) and SVMs (64.7±0.7%), the DNN performed slightly, but significantly better (Fig. 5H), suggesting that the nonlinear advantage of combining the features was not substantial here. In the case of ridge regression, the contribution of different features in predicting the emitter’s sex can be assessed (Fig. 5I), highlighting the USVs’ duration, volume and spectral breadth as the most distinguishing properties (compare also (Heckman et al. 2017)).

While the estimation of certain features is thus possible from the raw spectrograms, their predictiveness for the emitter’s sex stays comparatively low for both simple and advanced prediction algorithms. A sequential combination of the two networks would perform worse than either of the two. Hence, we hypothesize that the direct sex classification from spectrograms must rely on a different set of local or global features, not well captured in the present set of features.

## Discussion

The analysis of social communication in mice is a challenging task, which has been traction over the past years. In the present study we have built a set of cDNNs which are capable of classifying the emitter’s sex from single vocalizations with substantial accuracy. We find that this performance is dominated by sex-specific features, but individual differences in vocalization are also detectable and contribute to the overall performance. The cDNN classification vastly outperforms more traditional classifiers or DNNs trained on predefined features. Modern dimensional reduction techniques also fail to recover sex differences in vocalizations. Our results suggest that mice could be able to detect the sex of or even identify potential mating partners even from a distance on the basis of only a few vocalizations.

### Comparison with previous work on sex differences in vocalization

Previous work has approached the task of differentiating the sexes from the USVs using different techniques and different datasets. The results of the present study are particularly interesting, as a previous, yet very different analysis of the same general data set did not indicate a difference between the sexes (Hammerschmidt et al. 2012), The latter automatically calculated 8 acoustic features for each USV and applied clustering analysis with subsequent linear discriminant analysis. The analysis identified 3 clusters and the features contributing most to the discrimination were duration, frequency jumps and change. We find a similar mapping of features to clusters (not shown), however, both Hammerschmidt et al. 2012 and ourselves did not find differences relating to the emitter’s sex using these analysis techniques, i.e. both for dimensionality reduction and linear classification. Further, the present set of features also differed and was partly scored by humans, to allow a more reliable discovery of complex Gestalt-like features. While the human scoring was blinded and hence unbiased with respect to sex, it may have its own biases (e.g. non-constant criteria).

Previous studies on other data sets have lead to mixed results, with some studies finding differences in USV properties (Burke et al. 2018), others finding no structural differences but only on the level of vocalization rates (Guo and Holy 2007), However, to our knowledge, no study was so far able to detect differences on the level of single vocalizations, as in the present analysis.

### Contributions of sex-specific and individual properties

While the main focus of the present study was on identifying the sex of the emitter from single USVs, the results also demonstrate that individual differences in vocalization properties contribute to the identification of the sex. Previous research has been conflicting on this issue, with a recent synthesis proposed that suggests a limited capability for learning and individual differentiation (Arriaga and Jarvis 2013), From the perspective of classifying the sex, this is undesired, and we therefore checked to which degree the network takes individual properties into account when classifying sex. The resulting performance when testing the network only on entirely unused individual is lower (~78%), but it is well known that it will underestimate the true performance (since a powerful general estimator will tend to overfit the training set, e.g. (Sahani and Linden 2003)). Hence, if we expand our study to a larger set of mice, we predict that the sex-specific performance would increase to around ~80 (assuming that the performance is roughly between the performances for ‘individual properties used’ and ‘individual properties interfering’). More generally, however, the contribution of individual properties provides an interesting insight in its own right, regarding the developmental component of USV specialization, as the animals are otherwise genetically identical.

### Optimality of classification performance

The spectrotemporal structure of male and female USVs could be inherently overlapping to some degree and hence preclude perfect classification. How close the present classification comes to the best possible classification is not easy to assess. In human speech recognition, scores of speech intelligibility or other classifications can be readily obtained (using psychophysical experiments) to estimate the limits of performance, at least with respect to humans, often considered the gold standard. For comparison, human sex can be obtained with very high accuracy from vocalizations (DNN-based, 96.7%, (Buyukyilmaz and Cibikdiken 2016)), although we suspect classification was based on longer audio segments than those presently used. For mice, a behavioral experiment would have to be conducted, to estimate their limits of classifying the sex or identity of another mouse based on its USVs. Such an estimate would naturally be limited by the inherent variability of mouse behavior in the confines of an experimental setup. In addition, the present data did not include spatial tracking, hence, our classification may be hampered by the possibility that the animal switched between different types of interactions with the anesthetized conspecific.

### Biological versus laboratory relevance

Optimizing the chances of mating is essential for successfully passing on one’s genes. While there is agreement that one important function of mouse vocalizations is courtship (Guo and Holy 2007; Chabout et al. 2015; Heckman et al. 2016; Portfors and Perkel 2014) to which degree USVs contribute to the identification of a conspecific has not been fully resolved, partly due to the difficulty in attributing vocalizations to single animals during social interaction. Aside from the ethological value of deducing the sex from a vocalization, the present system also provides value for laboratory settings. The mouse strain used in the present setting is an often used laboratory strain, and in the context of social analysis the present set of analysis tools will be useful, in particular in combination with new, more refined tools for spatially attributing vocalizations to individual mice (Heckman et al. 2017)

### Generalization to other strains and social interactions

The present experimental dataset was restricted to a single strain in a single type of interaction, an awake male/female mouse with an anesthetized female mouse. It has been shown previously that different strains (Van Segbroeck et al. 2017; Sugimoto et al. 2011) and different social contexts (Chabout et al. 2015; Zala et al. 2017; Burke et al. 2018) influence the vocalization behavior of mice. As the present study depended partially on the availability of human classified USV features, we chose to work with a more limited set here. While not tested here, we predict that the performance of the current classifier would be reduced if directly applied to a larger set of strains and interactions. However, retraining a generalized version of the network, predicting both sex, strain and type of interaction would be straightforward and is planned as a follow-up study.

### Towards a complete analysis of social interactions in mice

The present set of classifiers provides an important building block for the quantitative analysis of social vocalizations in mice and other animals. However, there remain a number of generalizations to reach its full potential. Aside from adding additional strains and behavioral contexts, the extension to longer sequences of vocalizations is highly relevant. While we presently demonstrate that single vocalizations already provide a surprising amount of information about the sex and individual, we agree with previous studies that sequences of USVs play an important role in mouse communication (Holy and Guo 2005; Zakaria et al. 2012; Chabout et al. 2015) While neither of these studies differentiated between the sexes, we consider a combination of the two approaches an essential step of further automatizing and objectifying the analysis of mouse vocalizations.

## Acknowledgements

BE was supported by a Marie-Sklodowska Curie grant (#660328) and an NWO (ALWOP.346) grant during different parts of the project period. The authors would like to thank Hugo Huijs for manually classifying the entire set of vocalizations.

## Methods

We recorded vocalizations from mice and subsequently analyzed their structure to automatically classify the mouse’s sex and spectrotemporal properties from individual vocalizations. All experiments were performed with permission of the local authorities (Bezirksregierung Braunschweig) in accordance with the German Animal Protection Law. Data and analysis tools relevant to this publication will be made available in a data-sharing collection in the Donders data repository upon submission for publication (di.dcn.DSC_620840_0003_891(.

### Data Acquisition

The recordings were made from C57BL/6NCrl female and male mice older than 8 weeks, housed in groups of five in standard (Type II long) plastic cages, with food and water ad libitum. We used the resident-intruder paradigm to elicit ultrasonic vocalizations (USVs) from male and female ‘residents’. As intruders we used anaesthetized females to ensure that only the resident mouse could emit calls. Hence, vocalizations in a given recording could be uniquely assigned to an individual mouse with a given sex.

For the recording, resident mice (males and females) were first habituated to the room: Mice in their own home cage were placed on the desk in the recording room for 60 seconds. Subsequently, an unfamiliar intruder mouse was placed into the home cage of resident, and the vocalization behavior was recorded for 3 min using AVISOFT RECORDER 4.1. We recorded the USVs at a sampling frequency of 300 kHz. The microphone (UltraSoundGate CM16) was connected to a preamplifier (UltraSoundGate 116), which was connected to a computer (all sound recording hardware and software was from Avisoft Bioacoustics, Berlin, Germany). The recordings constitute a subset of the recordings collected in a previous study (Hammerschmidt et al. 2012)

Overall, 10055 vocalizations were recorded from 17 mice in 17 sessions. Mice were only recorded once, such that 9 female and 8 male mice contributed to the data set. Male mice produced 542±97 (Mean ± SEM) vocalizations, while female mice produced 636±43 calls over the recording period of 3 min.

### Data Processing and Extraction of Vocalizations

An automatized procedure was used to detect and extract vocalizations from the continuous sound recordings. The procedure was based on existing code (Holy and Guo 2005), but extended in several ways to optimize extraction of both simple and complex vocalizations. The quality of extraction was manually checked on a randomly chosen subset of the vocalizations. We here describe all essential steps of the extraction algorithm for completeness (the code is provided in the repository alongside this manuscript, see above).

The continuous recording was first high-pass filtered (4th order Butterworth filter, 25 kHz) before transforming it into a spectrogram (using the norm of the fast fourier transform for 500 samples (1.67 ms) windows using 50% consecutive window overlap, leading to an effective spectrogram sampling rate of ~1.2 kHz). Spectrograms were converted to a sparse representation by setting all values below 70th percentile to 0, which eliminated most close-to-silent background noise bins. Vocalizations were identified using a combination of multiple criteria (defined below), i.e. exceeding a certain spectral power, maintaining spectral continuity, and lie above a frequency threshold. Vocalizations that followed each other at intervals <15 ms were subsequently merged again and treated as a single vocalization. For later classification, each vocalization was represented as a 100×100 matrix, encoded as uint8. Hence, vocalizations longer than 100 ms (11.7%) were included shortened.

The three criteria above were defined as follows:

- *Spectral energy:* The distribution of spectral energies was computed across the entire spectrogram, and only time-point were kept in which any of the bins exceed 99.8% of the distribution (manually estimated). While this threshold seems high, we verified manually that this threshold did not exclude any clearly recognizable vocalizations. It reflects the relative rarity of bins containing energy from a vocalization.
- *Spectral continuity:* We tracked the maximum position across frequencies for each time-step, and computed the size of the difference in frequency location between time-steps. These differences were then accumulated, centered at each time-point for 3.3 ms (4 steps), in both directions. Their minimum was compared to a threshold of 15, which corresponds to ~3.4 kHz/ms. Hence, the vocalization was accepted if the central line did not change too much. The threshold was set generously, in order to also include more broadband vocalizations. Manual inspection indicated that no vocalizations were missed due to a too stringent spectral purity criterion.
- *Frequency threshold:* Despite the high-pass filtering some low-frequency environmental noises can contaminate higher frequency regions. We there excluded all vocalizations whos mean spectral energy was <25 kHz.

Only if these three properties were fulfilled, we included a segment of the data into the set of vocalizations. The segmentation code is included in the above repository (VocCollector, VocAnalyzer), with the parameters set as follows: SpecDiscThresh = 0.8, MergeClose = 0.015.

### Dimensionality Reduction

For initial exploratory analysis we performed dimensionality reduction on the human-classified features and the full spectrograms. Both principal component analysis (PCA) and t-statistic neighborhood embedding (tSNE, (Maaten and Hinton 2008), were applied to both datasets (see Fig. 2). tSNE was run in Matlab using the original implementation from (Maaten and Hinton 2008), using the following parameters (initial reduction using PCA to dimension 9 for features and 100 for spectrograms; perplexity: 30).

### Classification

We performed automatic classification of the emitter’s sex, as well as several properties of individual vocalizations (see below). While the sex was directly known, the ground truth for most of the properties was estimated by two independent human evaluators, who had no access to the emitting sex during scoring.

**Vocalization Properties** We defined a range of 10 properties which served to describe the shape of a vocalization.

- *Direction:* whether the vocalization is composed mostly flat, descending or ascending pieces, values are 0,-1,1, respectively
- *Peaks:* number of clearly identifiable peaks, values: integers ≥ 0.
- *Breaks:* number of times the main frequency whistle is disconnected, values: integers ≥ 0
- *Broadband:* whether the frequency content is narrow (0) or broadband (1). Values: [0,1]
- *Tremolo:* whether there is a sinusoidal variation on the main whistle frequency. Values: [0,1]
- *Complexity:* whether the overall vocalization is simple or complex. Values: [0,1]
- *Duration:* overall duration of the vocalization. Values > 0 ms
- *Frequency:* average frequency of the vocalization. Values > 0 Hz
- *Power*, average power of the vocalization. Values > 0 dB^2^

The first 6 properties were scored by the human evaluators, while the latter 3 properties were estimated automatically and added to the manually scored properties.

**Performance evaluation** The quality of classification was assessed using crossvalidation, i.e. by training on a subset of the data (90%) and evaluating the performance on a separate test set (remaining 10%). For significance assessment, we divided the overall set into 10 non-overlapping test sets (permutation draw, with their corresponding training sets), and performed 10 individual estimates. Performance was assessed as percent correct, i.e. the number of correct classifications in comparison with the total number of the same type. We performed another control, where the test set was just one recording session, which allowed us to verify that the classification was based on sex rather than individual (see Fig. 4B). Here, the size of the test set was determined by the number of vocalizations of each individual.

### Deep Neural Network Classification

For classification of sexes, features and individuals, we implemented several neural networks of standard architectures containing multiple convolutional and fully connected layers (cDNNs, see details below). The networks were implemented and trained in python 3.6 using the TensorFlow toolkit (Abadi et al. 2015) Estimations were run on single GPUs (GTX770, 4 GB and GTX1070, 8 GB).

The networks were trained using standard techniques, i.e. regularisation of parameters using batch-normalization (Ioffe and Szegedy 2015) and dropout (Srivastava et al. 2014) Batch normalization was applied for all network layers, both convolutional and fully connected. The sizes and the number of features for convolutional kernels were selected according to the parameters commonly used for natural images processing networks (specific architectural details are displayed in each figure). For training, stochastic optimization was performed using ADAM (Kingma and Ba 2014) The rectified linear unit (ReLU) was used as the activation function for all layers except for the output layer. For the output layer, the sigmoid activation function was used. The cross entropy loss function was used for minimization. The initial weight values of all layers were set using Xavier initialization (Xavier Glorot and Yoshua Bengio 2010)

*Spectrogram-to-Sex, Spectrogram-to-Individual and Spectrogram-to-Features Networks* In total, 6 convolutional networks taking spectrograms as their input were trained: *Spectrogram-to-Sex, Spectrogram-to-Individual* and 4 *Spectrogram-to-Features* networks including direction *(Spectrogram-to-Direction)*, number of peaks *(Spectrogram-to-Peaks)*, number of breaks *(Spectrogram-to-Breaks)* and broadband property *(Spectrogram-to-Broadband)* detection networks.

The networks consisted of 6 convolutional and 3 fully connected layers. The detailed layer properties are represented in the Table 1. The output layer of *Spectrogram-to-Sex* network contained single element, representing the detected sex (the probability of being male). The *Spectrogram-to-Direction, Spectrogram-to-Peaks and Spectrogram-to-Breaks* networks’ output layers contained the number of elements equal to the number of detected classes. Since only a few samples had number of peaks or breaks more than 3, the data set was restricted to 4 classes: no peaks(breaks), 1 peak(break), 2 peaks(breaks), 3 or more peaks(breaks). So, the output layers of *Spectrogram-to-Peaks and Spectrogram-to-Breaks* networks consisted of 4 elements representing each class. The *Spectrogram-to-Broadband* network output layer contained single element representing *Broadband* value.

**Table 1.** The architecture of Spectrogram-to-Sex and Spectrogram-to-Features networks

| Layer | Properties |
| --- | --- |
| Convolutional 1 | 10x10, 256 units, stride = 2 |
| Convolutional 2 | 5x5, 256 units, stride = 2 |
| Convolutional 3 | 3x3, 256 units, stride = 1 |
| Convolutional 4 | 3x3, 256 units, stride = 1 |
| Convolutional 5 | 3x3, 256 units, stride = 1 |
| Convolutional 6 | 3x3, 256 units, stride = 1 |
| Fully connected 1 | 120 (80*) units |
| Fully connected 2 | 120 (80*) units |
| Fully connected 3 | 120 (80*) units |

\* For Spectrogram-to-Broadband feature network we used 80 unit fully connected layers

Table 1. The architecture of Spectrogram-to-Sex and Spectrogram-to-Features networks

#### Features-to-Sex Network

The networks used for classification of sexes based on the spectrogram features consisted of 4 fully connected layers. The first layer received inputs from 9-dimensional vectors representing feature values: duration, average frequency, average volume, direction. number of peaks, number of breaks, “broadband” value, “tremolo” value, “complex” value. The output layer contained single element, representing the detected sex (the probability of being male). The detailed layer properties are represented in the Table 2.

**Table 2.** The architecture of Feature-to-Sex network

| Layer | Size |
| --- | --- |
| Fully connected 1 | 70 units |
| Fully connected 2 | 70 units |
| Fully connected 3 | 70 units |
| Fully connected 4 | 70 units |

Table 2. The architecture of Feature-to-Sex network

##### Input data preparation and augmentation

The original source spectrograms were represented as Nx233 images where N is rounded duration of the spectrogram in milliseconds. The values were in the range [0,255]. To represent the spectrograms as 100×100 images they were cut to 100 ms threshold and rescaled to Mx100 size keeping the original aspect ratio, M is the scaled thresholded duration. The rescaled matrix was aligned to the left side of the resulting image, the unused space was filled with gaussian noise with variance = 0.01.

For DNNs, we implemented on-the-fly data augmentation to enlarge the input dataset. For each image being fed to the network during training session a set of modifications were applied including start and end times clip (up to 10% percent of original duration), intensity (volume) amplification with a random coefficient drawn from the range [0.5, 1.5] and the addition of gaussian noise with the variance randomly drawn from the range [0, 0.01]. We used the scikit-image package (van der Walt et al. 2014) routines for implementing data augmentation operations.

Different approaches were used to compensate the asymmetry in occurrence of different classes (e.g. for there were 32% more female than male vocalizations), we used different procedures for the three classification methods: For Ridge and SVM, the number of male and female vocalizations was equalized in the training sets by enlarging the smaller set to the same size as the biggest set with copies randomly drawn from the same set. For DNNs, a loss-function was used with weighting according to the occurrence in the different classes.

##### Training protocols

For *Spectrogram-to-Sex, Spectrogram-to-Individual* and the training protocol included 3 stages with different training parameters set. (Table 3). For *Spectrogram-to-Features* and *Features-to-Gender* the training protocols included 2 stages (Table. 4 and 5). Batch size was set to 64 samples for all networks.

**Table 3.**
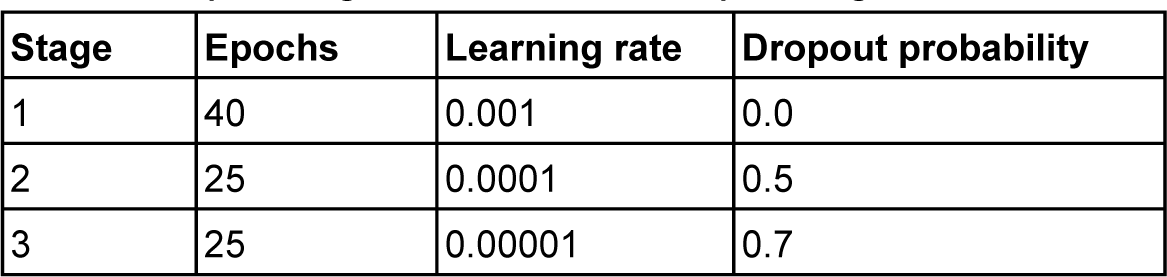
*Spectrogram-to-Sex* and *Spectrogram-to-Individual* networks training protocol

**Table 4.**
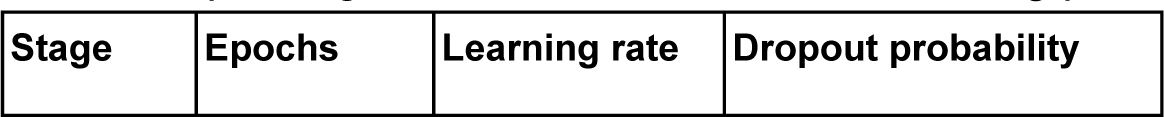
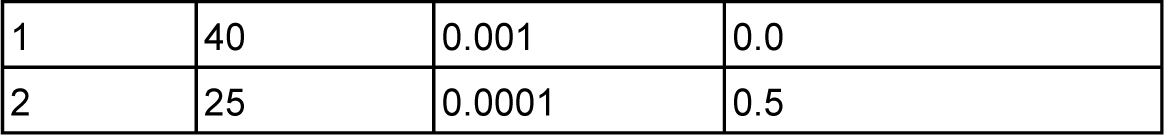
*Spectrogram-to-Features* networks training protocol

**Table 5.** *Features-to-Gender* network training protocol

| Stage | Epochs | Learning rate | Dropout probability |
| --- | --- | --- | --- |
| 1 | 700 | 0.001 | 0.0 |
| 2 | 400 | 0.0001 | 0.5 |

### Linear Regression and Support Vector Machines

To assess the contribution of basic linear prediction to the classification performance, we performed regularized (*r*(*idge*) regression by direct implementation of the normal equations in MATLAB. The performance was generally close to chance and did not depend much on the regularization parameter.

To check whether simple nonlinearities in the input space could account for the classification performance of the DNN we used support vector machine classification (Ben-Hur et al. 2008; Cortes and Vapnik 1995) using their implementation in MATLAB *(svmtrain, svmclassify).* We used the quadratic kernel for the spectrogram-based classification, and the linear (dot-product) kernel for the feature-based classification. The performance was above chance, however, much poorer than for DNN classification.

For both methods the identical crossvalidation sets were used as for the DNN estimation (see Fig. 3 and 5 for comparative results).

### Statistical Analysis

Generally, nonparametric tests were used to avoid distributional assumptions, i.e. Wilcoxon’s rank sum test for two group comparisons, and Kruskal-Wallis for single factor analysis of variance. When data were normally distributed, we checked that statistical conclusions were the same for the corresponding test, i.e. t-test or ANOVA, respectively. Effect sizes were computed as the variance accounted by a given factor, divided by the total variance. Error bars indicate 1 SEM (standard error of the mean). Post-hoc pairwise multiple comparisons were assessed using Bonferroni correction. All statistical analyses were performed using the statistics toolbox in MATLAB (The Mathworks, Natick).

